# The high energetic cost of rapid force development in cyclic muscle contraction

**DOI:** 10.1101/2020.08.25.266965

**Authors:** Tim J. van der Zee, Arthur D. Kuo

## Abstract

Muscles consume metabolic energy for active movement, particularly when performing mechanical work or producing force. Less appreciated is the cost for activating and deactivating muscle quickly, which adds considerably to the overall cost of cyclic force production (Chasiotis et al., 1987). But the cost relative to mechanical work, which features in many movements, is unknown. We therefore tested whether fast activation-deactivation is costly compared to performing work or producing isometric force. We hypothesized that metabolic cost would increase with a proposed measure termed *force-rate* (rate of increase in muscle force) in cyclic tasks, separate from mechanical work or average force level. We tested humans (N = 9) producing cyclic knee extension torque against an isometric dynamometer (torque 22 N-m, cyclic waveform frequencies 0.5 – 2.5 Hz), while also quantifying the force and work of muscle fascicles against series elasticity (with ultrasonography), along with metabolic rate through respirometry. Net metabolic rate increased by more than fourfold (10.5 to 46.7 W) with waveform frequency. At high frequencies, the hypothesized force-rate cost accounted for nearly half (41%) of energy expenditure. This exceeded the cost for average force (17%) and was comparable to the cost for shortening work (42%). The energetic cost is explained by a simple first-order model of rate-limiting steps in muscle contraction, primarily crossbridge dynamics. The force-rate cost could contribute substantially to the overall cost of movements that require cyclic muscle activation, such as locomotion.

**Summary statement:** The energetic cost of isometric muscle force production during cyclic muscle contraction increases sharply with cycle frequency and in proportion to the rate of force development

## Introduction

Humans often expend energy to perform movement tasks where muscles are only intermittently or cyclically active. Two notable contributions to the metabolic cost of such tasks are for the mechanical work performed by muscle fascicles (Abbott et al., 1952; Margaria, 1968), and for the force exerted when fascicles are isometric (Crow & Kushmerick, 1982). Less appreciated is the cost for muscle activation and deactivation during cyclic conditions. This cost increases with activation frequency (Hogan et al., 1998) or the rate of force production (Doke & Kuo, 2007), and can equal or even exceed the cost for producing continuous isometric force (Chasiotis et al., 1987). However, there is also substantial metabolic energy expended in tasks that also entail work, for example locomotion with the lower extremity (Margaria, 1976) or reaching with the upper extremity (H. J. Huang et al., 2012). But the relative cost for activating muscle vs. performing work remains unknown. It is therefore helpful to determine whether the rate of muscle force production has a cost comparable to work and force production.

There is clearly an energetic cost for activating muscle under intermittent, isometric conditions. This has been demonstrated with square-wave, on-off activation patterns, which yield a considerably higher metabolic cost than continuous activation at similar overall contraction duration (Chasiotis et al., 1987; Spriet et al., 1988). Intermittent contraction also requires more metabolic energy when contraction frequency is higher (Bergström & Hultman, 1988; Hogan et al., 1998) or when duration of active force production is smaller (Beck et al., 2020), even when accounting for the cost of maintaining isometric force, which is roughly proportional to the force-time integral (Crow & Kushmerick, 1982). Instead, the metabolic cost may be related to the frequency of muscle activation-deactivation (Bergström & Hultman, 1988), leading to a doubling of energetic cost per unit force compared to continuous isometric force (Chasiotis et al., 1987). The underlying mechanism may be different than for doing work, as the activation-deactivation cost has been attributed to active calcium transport rather than to crossbridge cycling (Hogan et al., 1998).

There is also a similar metabolic cost for cyclic, non-isometric movements. For example, cyclic leg swinging (Doke et al., 2005; Doke & Kuo, 2007) and ankle bouncing (Dean & Kuo, 2011) have costs increasing markedly with movement frequency (e.g., three-to four-fold for a 45% increase in frequency; Doke et al., 2005), and not explained by the amount of mechanical work (Doke & Kuo, 2007). This cost has instead been related to the rate of force production, or *force-rate*, which may be considered an analogue of the square-wave frequency, but for continuous, non-isometric movements. The cost of leg swinging, including a force-rate component, could account for one-third of the net metabolic cost of walking (Doke et al., 2005). The hypothesized force-rate cost may therefore be relevant to typical human activities.

However, it remains unclear to what extent the force-rate cost contributes to overall metabolic cost of movement. Everyday movements can include some combination of muscle mechanical work, force production, and force-rate costs. But the previous studies have either eliminated mechanical work with isometric conditions (e.g., Chasiotis et al., 1987), or controlled for it without directly estimating the amount of work performed (e.g., Doke & Kuo, 2007). This makes it difficult to separate the force-rate cost and quantify its contribution relative to both work and force. Another confounding factor is series elasticity, which stores and returns elastic energy not observable from the work of the overall muscle-tendon unit. Even in apparently isometric conditions, muscle fascicles may perform shortening work against series elastic tissues, with a meaningful contribution to overall cost. After all, muscle fascicle work is biochemically and thermodynamically constrained to cost metabolic energy (Barclay, 2015). The hypothesized cost of force-rate therefore needs to be quantified alongside the actual shortening work performed by fascicles.

The current study is intended to address that gap, by testing for a metabolic cost of cyclic muscle contraction, while accounting for and estimating the costs for mechanical work and force. We simultaneously quantified work (against series elasticity), force, and force-rate during a cyclic force production task, along with the overall metabolic energy expenditure. The task was to cyclically produce voluntary knee extension torque against an isometric dynamometer, at an amplitude and range of frequencies comparable to everyday human movements. We hypothesized that (1) work and force-time integral fall short in accounting for metabolic cost at higher frequencies, and (2) that the surplus cost is related to muscle force-rate. We tested this by parametrically relating each contribution as a function of waveform frequency and force-rate, and testing whether force-rate is separable from and comparable to the costs of work and force-time integral.

## Methods

We estimated quadriceps muscle force and mechanical work, and net metabolic rate as healthy adults produced cyclic, isometric knee extension torque (Fig. 1). Increasing frequency of cyclic torque was expected to yield greater fluctuations in muscle force. We conducted a set of energetics trials to test for an associated metabolic cost, and a set of mechanics trials to quantify muscle fascicle mechanical work (against series elasticity) and force via ultrasound imaging.

**Fig. 1:**
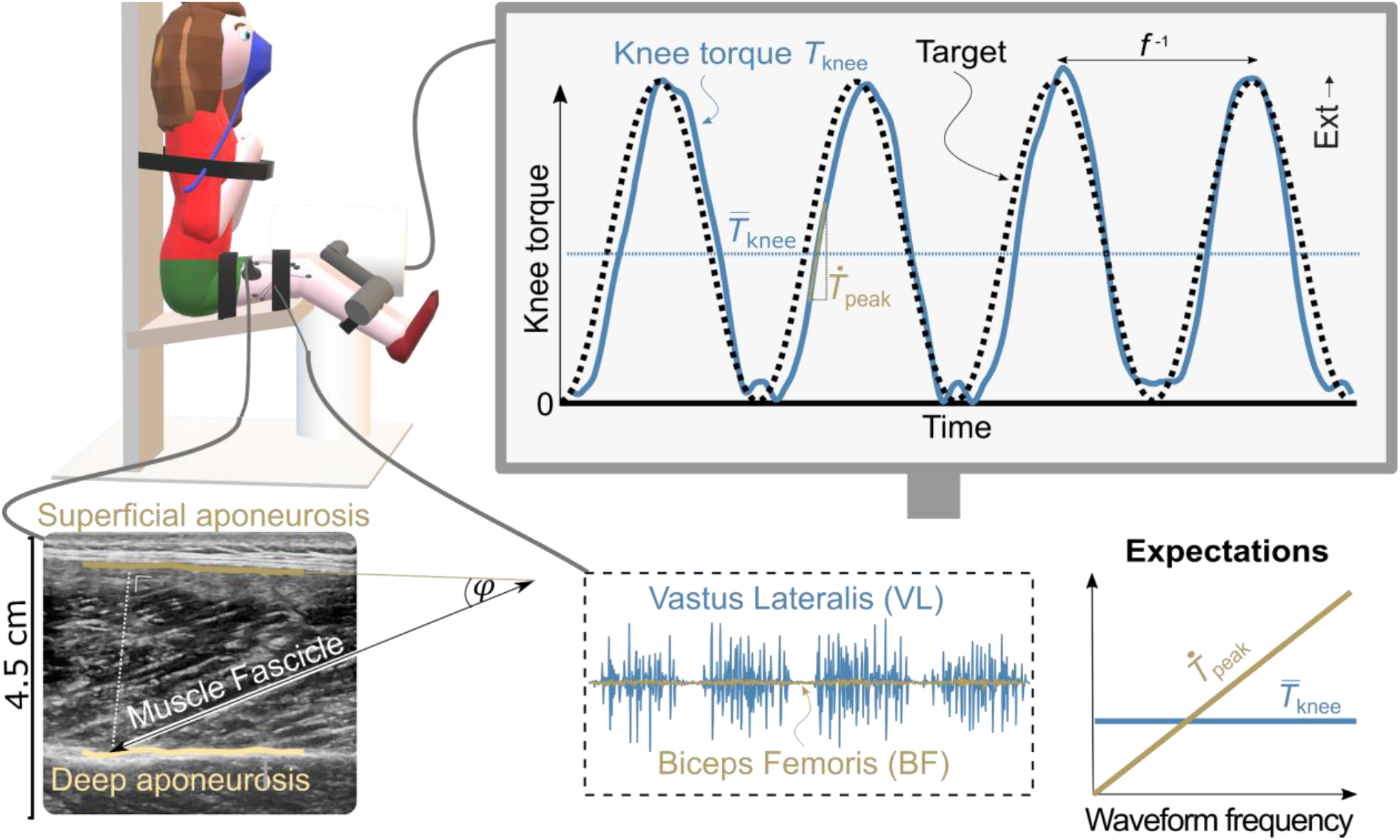
Experimental set-up for estimating metabolic cost of cyclic, isometric torque production. Participants produced knee extension torque to match a displayed waveform target, while respirometry, electromyography (EMG) and ultrasound were recorded. Muscle fascicle length and pennation angle were estimated from ultrasound, and combined with dynamometer to estimate fascicle force and work (against series elasticity). Sinusoidal torques of increasing waveform frequency *f* (and fixed amplitude) were expected to result in increasing peak torque-rate 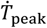 and fixed mean knee torque 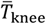, and therefore increasing fascicle force-rate and energetic cost.

Nine healthy participants (6 male, 3 female, leg length = 87.4 ± 4.9 cm, body mass = 70.6 ± 13.1 kg, mean ± s.d.) produced cyclic bilateral knee extension torque against an isometric dynamometer. The task was intended to require about 10% of maximal quadriceps muscle force and to impose contraction frequencies *f* within ranges encountered in daily life (mean 11 N-m and 0.5-2.5 Hz, respectively). Real-time visual feedback was displayed to participants, showing measured and target waveforms (Fig. 1) for each frequency, all at a fixed torque range (0 - 22 N-m) from combined legs. Torques were exerted at 20 deg knee flexion (relative to straight leg) and measured with a dynamometer chair (Biodex, Biodex Medical Systems, NY, USA), which was also used to strap and constrain movement of the trunk and legs. As baseline, we evaluated knee extension torque during maximum voluntary contraction (MVC) at 20 deg knee flexion (97 ± 20 N-m). As a reference, we also evaluated MVC extension torques at 70 deg knee flexion (221 ± 70 N-m), closer to the plateau of the knee extension torque-angle relationship (e.g. Kulig et al., 1984). For each condition, we quantified knee torque in terms of its time average 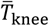, peak amplitude, and peak torque-rate 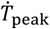 (Fig. 1). Peak torque-rate 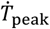 was estimated from the change in torque from 50% to 150% of average, divided by the time duration of that change. Knee torque *T*_knee_ and its peak rate 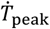 were then divided by estimated muscle moment arm to estimate muscle fascicle force *F*_fas_ and its peak rate 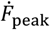. Prior to data collection, participants provided their written informed consent according to University of Calgary procedures.

The energetics trials entailed cyclic torque production while respirometry data were recorded. Participants were first given practice with matching real-time torque targets for at least 5 min. They then performed the five experimental task conditions for 6 min each, with torque and respirometry (rates of oxygen consumption 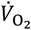 and carbon dioxide production 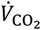; K5 system, Cosmed, Rome) averaged only for the final 3 min, to allow time to reach steady-state activity. Metabolic rate was determined from 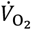 and the ratio between 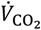 and 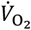 rates (Lusk, 1909). We also recorded electromyography (EMG) to monitor muscle activity and co-contraction (vastus lateralis, rectus femoris, biceps femoris, abbreviated as VL, RF and BF respectively). EMG amplitude was determined from the band-pass filtered recordings (cut-off frequency *f*_*c*_ = 30-500 Hz, Butterworth) and low-pass filtered to obtain the envelop (*f*_*c*_ = 5 Hz, Butterworth) (Hof, 1984). EMG amplitude was defined as the absolute of the Hilbert transformed signal (N. E. Huang et al., 1998), expressed relative to its maximum value during MVC and corrected for background noise using baseline-subtraction (La Delfa et al., 2014). The order of task conditions was randomized, and with at least 1 min of rest between conditions.

The separate mechanics trials employed ultrasound imaging to estimate muscle fascicle shortening and work against series elasticity. These trials consisted of single-legged isometric torque production at three knee angles (*θ*_knee_ = 15, 20 and 25 deg), guided by visual feedback of a low frequency triangle waveform torque target (*f* = 0.02 Hz, range = 0-16 N-m). Images were obtained for VL using ultrasound (5 cm probe, 11 MHz basic-mode; Logiq E9, General Electric, Fairfield, USA), recorded at 30 Hz and approximately synchronized with torque recordings (via a sync pulse). Separate trials were performed for each of both legs and then averaged between legs.

Muscle fascicle lengths *L*_fas_ and muscle pennation angles *φ* were estimated using a custom ultrasound algorithm (van der Zee & Kuo, 2020). The SEE length changes Δ*L*_SEE_ were estimated from changes in fascicle lengths *L*_fas_ (sampled as a function of *T*_knee_ at 450 uniformly spaced points) and pennation angles *φ* (Fukunaga et al., 2001) and were assumed to be solely dependent on SEE force *F*_SEE_. Consequently, at fixed knee torque *T*_knee_ (and thus fixed *F*_SEE_), quadriceps muscle-tendon complex (MTC) length change could be estimated from the component of fascicle length change along the SEE. The moment arm *r*_MTC_ of the quadriceps MTC about the knee (at *θ*_knee_ = 20 deg) was estimated from the average difference in MTC length between the 15 deg and the 25 deg trials, divided by the corresponding knee angle difference (i.e. Δ*θ*_knee_ = 10 deg). The corresponding force *F*_SEE_ was estimated as knee torque *T*_knee_ divided by *r*_MTC_.

The effects of knee torque *T*_knee_ on muscle fascicle length change Δ*L*_fas_ and pennation angle change Δ*φ*, as well as the effect of SEE force *F*_SEE_ on SEE length change Δ*L*_SEE_ were averaged across knee angles and fitted using exponentials. An exponential toe region was used for the SEE force-length curve, appropriate for the relatively low torque levels, less than 10% of maximal torque (Kulig et al., 1984; Lichtwark & Wilson, 2008). The mechanical work done on SEE by the fascicle (*W*_fas_) was quantified by the integral of the (fitted) SEE force *F*_SEE_ with respect to the (fitted) SEE length *L*_SEE_, for the torques applied during the energetics trials. Fits were done for each subject individually; fascicle work estimates were averaged across subjects.

We hypothesized that overall metabolic cost should include contributions from muscle force-rate 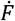, as well as from mechanical work and steady force production. The force-rate cost was hypothesized to be a consequence of rate-limiting dynamics between muscle excitation and force production (excitation-contraction dynamics, Fig. 2A), particularly due to the rate of cross-bridge binding (Brenner & Eisenberg, 1986). It should be noted that Hill-type muscle models (Zajac, 1989) do not generally include cross-bridge binding dynamics between calcium release (Ebashi and Endo, 1968) and force production. Here we illustrate the consequence of rate-limited force production with simple first-order dynamics, which show how an increasing amount of calcium would need to be released (for example via motor unit recruitment) to produce cyclic force waveforms of constant amplitude but increasing frequency (Fig. 2B). Calcium release requires active calcium pumping to deactivate muscle, with an associated metabolic cost *E*_FR_ increasing with rate of force production (FR for force-rate). This general concept is illustrated with an example time constant of 30 ms (Fig. 2B).

**Fig. 2:**
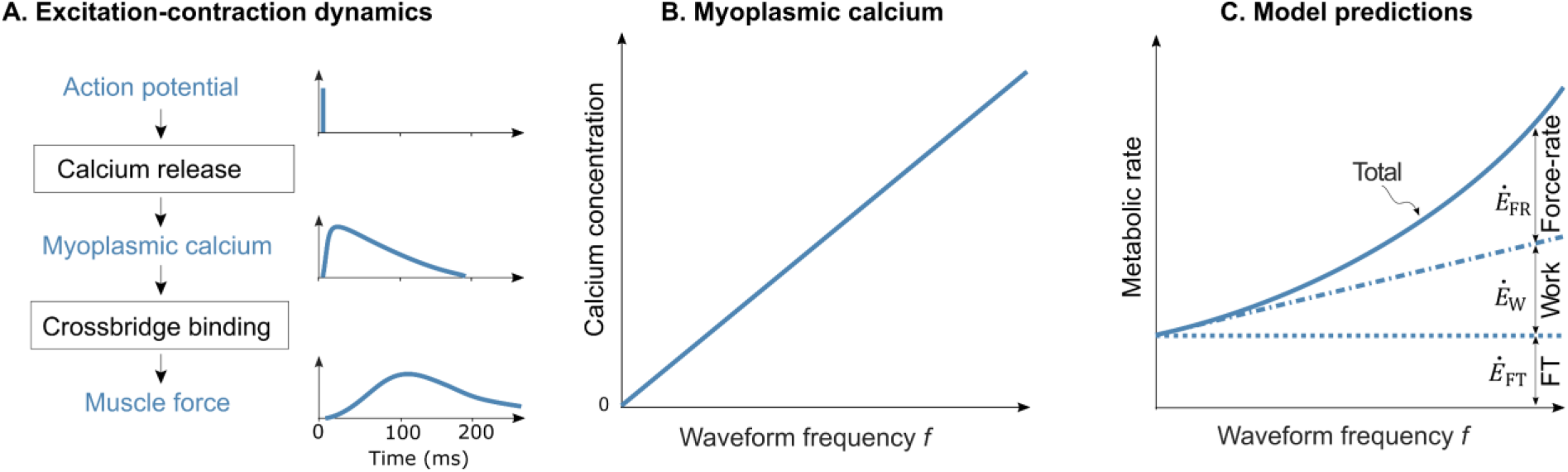
Hypothesized mechanism for force-rate cost. (A) Excitation-contraction dynamics start with an action potential that triggers calcium release into the myoplasm, which enables cross-bridge binding and muscle force production. The rate-limiting effect of cross-bridge binding is illustrated by hypothetical transients of action potential, myoplasmic calcium, and muscle force vs. time. (B) The excitation-contraction dynamics require increasingly large input (myoplasmic calcium) amplitudes, for cyclic forces of increasing waveform frequency and constant amplitude. As demonstration, this is modeled as a first-order dynamical system (*τ* = 30 ms), with greater calcium (e.g. via recruitment of additional motor units) costing energy for active transport following force production. (C) This results in greater metabolic cost per muscle contraction, as well as per time. Associated metabolic rate *Ė*_FR_ for force-rate (FR) is expected to increase quadratically with waveform frequency. In contrast, metabolic rate for force maintenance *Ė*_FT_ and mechanical work *Ė*_w_ (FT for force-time integral, W for work) are expected to remain fixed and increase linearly, respectively.

The force-rate cost hypothesis may be expressed as a metabolic cost per contraction, and experimentally tested in terms of a metabolic rate (or cost per time). The cost per muscle contraction *E*_FR_ should increase in proportion to how quickly the force *F* increases,

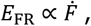

and particularly the peak force-rate 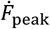. For a sinusoidal force waveform of fixed amplitude, peak force-rate is expected to increase with waveform frequency *f*,

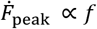

Under steady-state conditions, the corresponding metabolic rate *Ė*_FR_ (Fig. 2C) depends on the cost per contraction *E*_FR_ and on the frequency of contractions, yielding

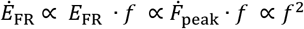

The work and force-time contributions should have separable dependencies on frequency *f*. Despite an isometric joint, muscle fascicles perform work, against series elastic tendon and aponeurosis, and imperfectly rigid leg and dynamometer. For a fixed torque amplitude, work should be performed in fixed amount per contraction, and the rate of mechanical work 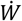 should therefore increase with the rate of contractions, or waveform frequency *f*. Assuming fixed biochemical costs for fascicle work, the corresponding metabolic rate *Ė*_*W*_ for work production should increase as

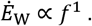

The force maintenance contribution is for energy expended even when muscle fascicles are isometric. Metabolic cost should be proportional to the force amplitude and duration of force (or *force-time integral*, Crow & Kushmerick, 1982). Due to constant amplitude of the waveform, average knee torque 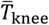 and therefore average muscle force was expected to be independent of waveform frequency. The corresponding force-time metabolic rate *Ė*_FT_ should be independent of waveform frequency,

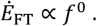

In experiment, we tested for an overall metabolic cost with all three of these contributions. We hypothesized that net metabolic rate *Ė*_net_ would be the sum of force-rate, work, and force-time integral terms:

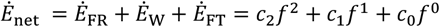

where “net” is defined as gross minus the cost of quiet sitting. The expected cost of performing fascicle work is derived from the inverse product of the efficiencies of (1) obtaining ATP from food stuff (about 60%, van Ingen Schenau et al., 1997) and (2) cross-bridge formation (about 50%, Barclay, 2015), yielding the linear coefficient *c*_1_ of 3.33. The other coefficients (*c*_0_ and *c*_2_) were determined using regression with *f* as independent variable. Unless stated otherwise, standard deviations (s.d.) refer to between-subjects variability.

## Results

Prior to examining the metabolic rate of cyclic force production, we first examine the experimental conditions, which showed that participants matched target torques well. As expected, participants knee torque *T*_knee_(*t*) had approximately sinusoidal waveform, with constant average 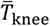 and variable peak rate 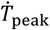. There was no significant dependency of waveform frequency *f* on average torque 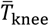 (slope 0.2±0.4 N-m·Hz^-1^, mean ± 95% confidence interval CI, *P =* 0.5, linear regression). Average knee torque 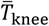 and torque minima and maxima were 10.7±1.0 N-m, 1.0±1.0 N-m and 21.8±1.2 N-m respectively (mean ± s.d. across subjects and conditions; Fig. 3A). Despite constant average torque 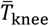, quadriceps EMG amplitude increased with waveform frequency by about 76% (VL: 2.1±0.5 %MVC at 0.5 Hz to 3.7±0.6% MVC at 2.5 Hz, mean ± s.d.). The increase was approximately linear in frequency *f*, at rates of 0.8±0.2 %MVC·Hz^-1^, 1.0±0.3 %MVC·Hz^-1^ and 0.2±0.1 %MVC·Hz^-1^ for VL, RF and BF respectively (mean ± 95% CI, repeated measures linear regression). There was some co-contraction, with BF EMG amplitude less than 1% MVC in all conditions (Fig. 3C), on average about one-eighth of VL EMG amplitudes. Peak knee torque-rate 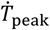 increased with waveform frequency *f* (slope 68.3±3.4 N-m, mean ± 95% confidence interval CI, linear regression without offset), and closely resembled the increase expected from the torque targets (slope 69.1 N-m, *R*^2^ *=* 0.84, RMSE = 1.1 N-m, root-mean-square-error; Fig. 3B).

**Fig. 3:**
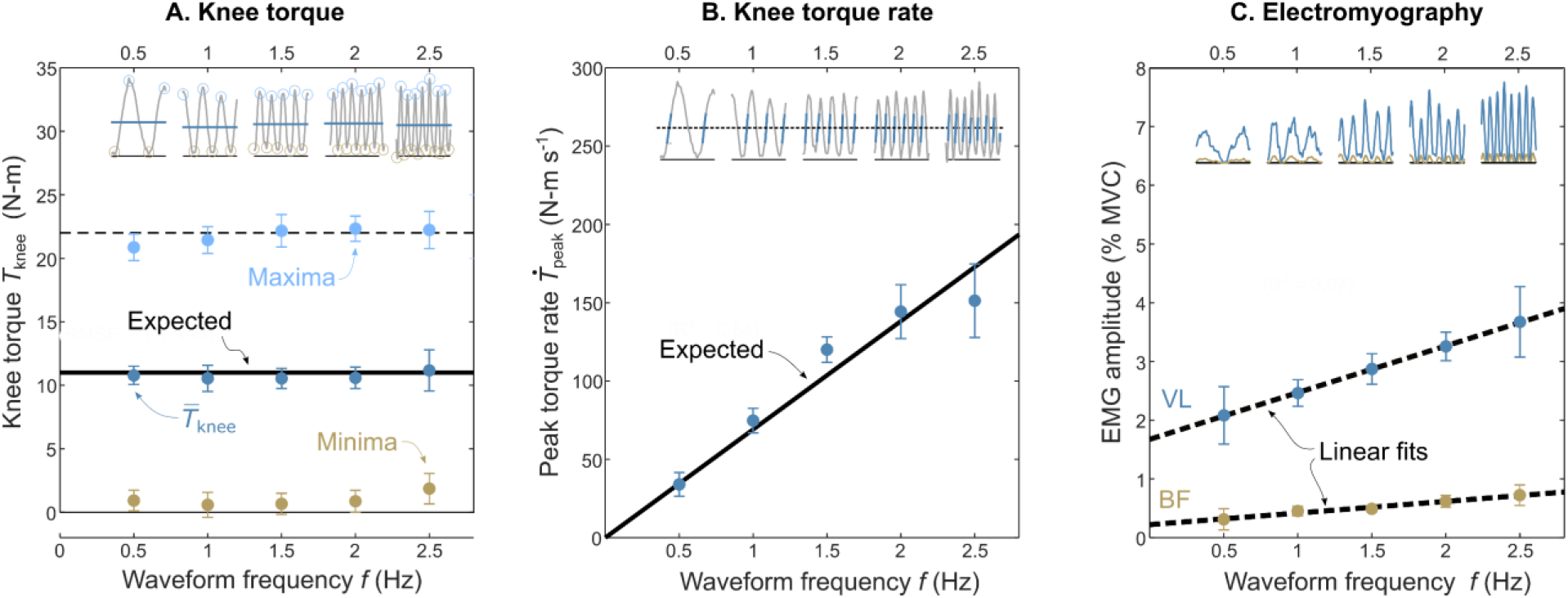
Experimentally measured torque, torque-rate, and electromyography vs. waveform frequency. (A) Knee torque vs. waveform frequency *f*, in terms of time-average, minima, and maxima of waveforms (shown in insets). Time-average torque 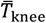 was relatively constant across conditions, comparable to the expected 11 N-m. (B) Peak torque-rate vs. waveform frequency *f*. Torque-rate 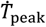 (shown in inset) increased with frequency *f*, comparable to expected (solid line). (C) Electromyography (EMG) amplitudes vs. frequency *f*, for Vastus Lateralis (VL) and Biceps Femoris (BF). EMG amplitude increased approximately linearly with waveform frequency. Filled symbols denote means across subjects (*N =* 9); error bars denote s.d.

Muscle fascicles performed work during isometric torque production, as evidenced by fascicle shortening (Fig. 4A) and pennation angle changes (Fig. 4B) observed during ultrasound trials. As a baseline, vastus lateralis fascicle length was estimated at 9.5±0.8 cm (mean ± s.d.) during rest (20 deg knee flexion). Fascicle lengths decreased with greater torque, as described by a fitted exponential (R^2^ = 0.38; see Fig. 4A). At 11 N-m, muscle fascicles had shortened 1.3±0.6 cm. Pennation angle was 15.9±1.2 deg (mean ± s.d.) during rest. Pennation angle increased with greater torque, also described by a fitted exponential (R^2^ = 0.46; Fig. 4B). Pennation angle was 15.9±1.2 deg (mean ± s.d.) at rest, and increased by 3.5±1.3 deg at 11 N-m. Combining fascicle length and pennation angle changes with knee angle, quadriceps muscle-tendon complex moment arm was estimated at 5.2±2.3 cm (mean ± s.d.). These data also yielded an estimate of length change of series elastic elements, fit by an exponential toe region (R^2^ = 0.42; Fig. 4C). From that fit, muscle fascicle mechanical work *W*_fas_ was estimated at 1.2±0.6 J (mean ± s.d.) for an 11 N-m contraction. The increase in mechanical work rate 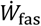 with waveform frequency was therefore about 2.4±1.2 J.

**Fig. 4:**
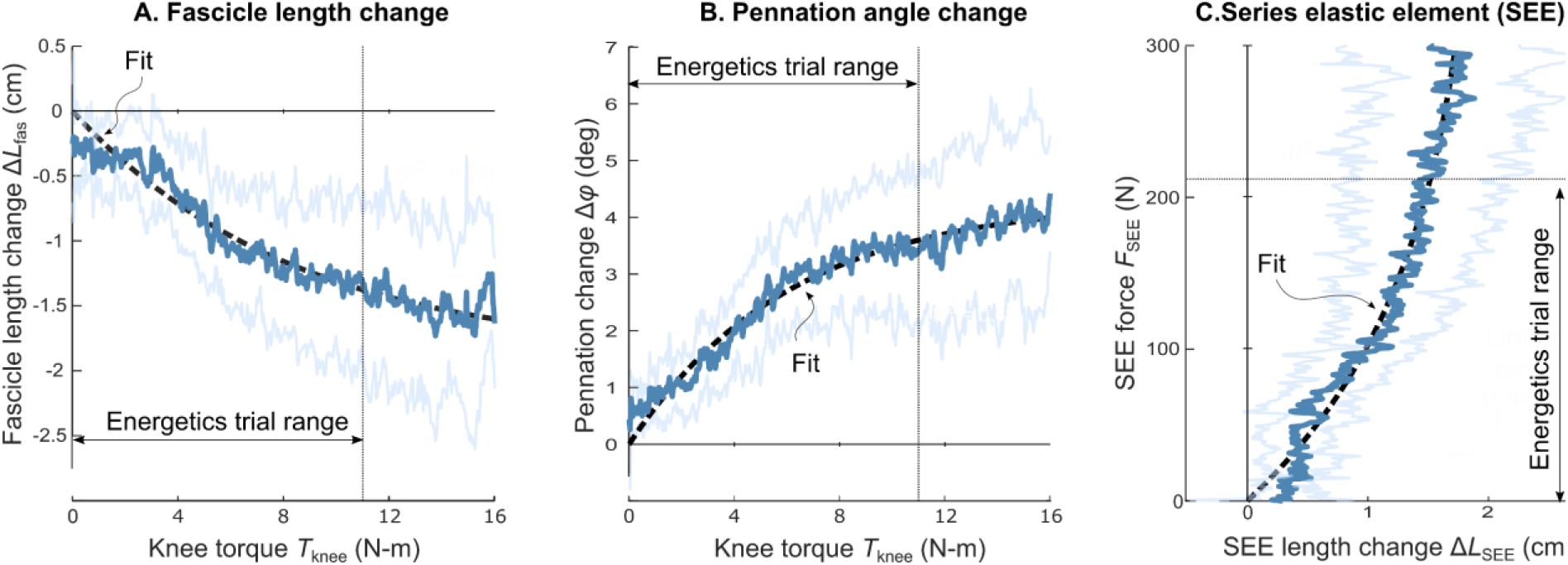
Muscle fascicle and series elastic element elongation as a function of knee torque. (A) Muscle fascicle length change (relative to relaxed) Δ*L*_*FAS*_ vs. knee torque *T*_knee_. Fascicle length decreased with greater knee torque, with exponential fit. (B) Muscle pennation angle change Δ*φ* vs. knee torque torque *T*_knee_. Pennation angle increased with greater knee torque, with exponential fit. (C) Series elastic element (SEE) force *F*_SEE_ vs. length change Δ*L*_SEE_. SEE length increased with greater SEE force, with fit to exponential toe region. Data here were obtained during isometric mechanics trials. Resulting fit was used to estimate mechanical work done by fascicles on SEEs during the energetics trials. Dark lines indicate means across subjects (*N =* 9); light lines denote s.d.

Net metabolic rate *Ė*_*net*_ increased with waveform frequency *f* (Fig. 5A), in agreement with predictions (*P* = 4·10^−6^, repeated measures regression). Net metabolic rate *Ė* increased from a value of 10.5 ± 13.2 (mean ± s.d.) at the lowest frequency (*f* = 0.5 Hz), to 46.8±8.4 W (mean ± s.d.) at the highest frequency (*f* = 2.5 Hz). The overall cost appeared to agree with hypothesized contributions from force-rate *Ė*_FR_, work *Ė*_W,_ and force-time *Ė*_FT_. At the highest frequency, these three costs were estimated at 19.1 W, 19.7 W and 7.9 W respectively, or about 41%, 42% and 17% of net metabolic rate *Ė*. Costs of force-rate *Ė*_FR_ and force-time *Ė*_FT_ had coefficients of 3.2±1.4 J·s and 7.9±10.2 W respectively (mean ± CI, *P* = 0.007, repeated measures regression). Examining the force-rate cost as a cost per contraction *E*_FR_, it increased approximately linearly with measured peak force-rate 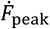 (2.6±1.9 · 10^−3^ m·s, mean ± CI, linear regression without offset; Fig. 5B).

**Fig. 5:**
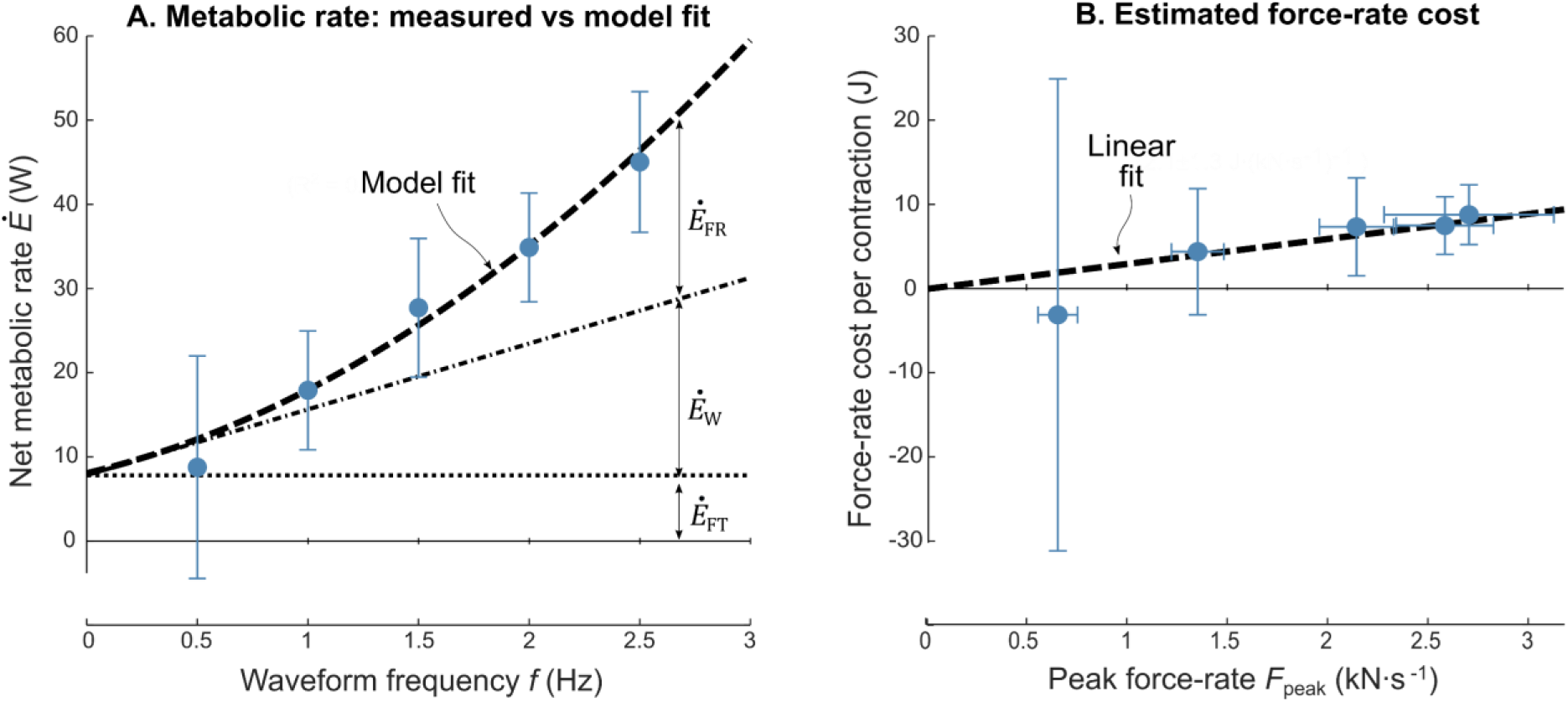
Metabolic cost of cyclic force production. (A) Net metabolic rate *Ė*_*net*_ versus waveform frequency *f*, along with three separate hypothesized contributing terms: (1) a force-time term *Ė*_FT_ independent of *f*, (2) a work term *Ė*_*W*_ increasing linearly with *f*, and (3) the hypothesized force-rate term *Ė*_FR_, increasing with *f*^2^. Net metabolic rate *Ė*_*net*_ was largely explained by a quadratic fit (R^2^ = 0.66) where the linear coefficient was fixed to the cost of work *c*_1_. (B) Force-rate cost per contraction *E*_FR_ versus measured peak force-rate 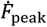. Force-rate cost *E*_FR_ (isolated from force-time and work terms) increased approximately linearly with measured peak force-rate 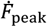. Filled symbols denote means across subjects (*N =* 9); error bars denote s.d.

## Discussion

The current study aimed at investigating the metabolic cost related to muscle force-rate, separately from both force and mechanical work costs. Net metabolic rate increased with the frequency of isometric torque production, faster than would be expected for force production alone. We also estimated muscle fascicle shortening, and observed non-negligible work performed by fascicles, but also insufficient to explain metabolic cost. A considerable portion of this cost can be ascribed to the proposed force-rate cost. We next examine the hypothesis in relation to the data, the potential mechanisms for metabolic cost, and implications for movement.

We first consider why force and work production do not explain the observed metabolic cost. The cyclic, isometric task resulted in nearly the same average force across conditions. The fascicle force-time integral was therefore unchanged as a function of frequency. Fascicles did, however, shorten and perform work that increased with frequency. But the expected metabolic cost for work *Ė*_*W*_, based on physiological (Margaria, 1968) and biochemical (Barclay, 2015) grounds, was far below the observed metabolic cost. Our estimate does not include the dependency of work coefficient *c*_1_ on contraction velocity, as described by a parabolic efficiency-velocity curve (Barclay, 2015). Mechanical efficiency is highest at about 33% of maximal contraction velocity (37 cm·s^-1^ for VL) (Barclay et al., 1993; Ruiter et al., 2000; van Soest & Bobbert, 1993), which is substantially higher than contraction velocity in the highest frequency condition here (6.5 cm·s^-1^). Therefore, higher waveform frequencies could allow for a higher mechanical efficiency and thereby a lower work cost, potentially making ours an overestimate of the work cost at higher frequencies. We have also treated force-time and work costs as separate and additive, when both result from the same mechanism, namely actomyosin ATPase activity (Barclay et al., 2007). Crossbridges have a fixed amount of energy per cycle, with much of it dissipated as heat in isometric conditions, as described by force-time coefficient *c*_0_. But that coefficient might be expected to decrease with contraction velocity, so that actomyosin ATPase activity might produce relatively more shortening work at higher contraction velocities. This would make the force-time cost an overestimate at higher frequencies. Yet another concern is muscle co-contraction, which could lead to greater cost not explained by shortening work. We did observe antagonist EMG of relatively small and linearly increasing amplitude, but consider it unlikely to explain the quadratically increasing energetic cost with waveform frequency. Most of the assumptions we have applied would lead to overestimation of work and force-time costs, so that their sum might in reality explain less than the 59% of net metabolic cost we have estimated.

This leaves a substantial cost explained by the hypothesized force-rate cost. The sharp increase in metabolic cost with muscle contraction frequency (Fig. 5) was consistent with the expected force-rate cost increasing with the square of contraction frequency. The hypothesis is based on the rate-limited dynamics between muscle excitation and force production, which acts as a low-pass filter of excitation and associated calcium release into the myoplasm. In the present experiment, increasing contraction frequencies called for higher force-rates (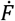, Fig. 3), which are expected to require more calcium release per muscle contraction. Active calcium pumping returns the calcium to the sarcoplasmic reticulum, with an associated metabolic cost (SERCA ATPase; (Inoue et al., 2019)). This contrasts with the energetic costs for force maintenance or work production, both associated with cycling of actomyosin crossbridges (actomyosin ATPase). Our hypothesis is that the combination of rate-limiting dynamics and calcium pumping imply a separate force-rate cost. This cost is in addition to the traditional force- and work-related costs, and could explain 41% (or more, if work and force-time costs are overestimated) of metabolic cost observed at the higher movement frequencies of the present experiment. In movement conditions that require cyclic muscle contraction, the force-rate cost could be quite substantial.

A source of uncertainty was our estimate of muscle shortening work. We estimated Vastus Lateralis (VL) muscle fascicle work (about 1.2 J, see Fig. 4) done while shortening against SEEs during isometric torque production. Whereas such shortening is similar as reported elsewhere (Ichinose et al., 1997), others have estimated considerably less shortening (e.g. 0.64 cm for VL) (van Soest & Bobbert, 1993). Some of the fascicle shortening we observed could be due to dynamometer deformation and tendon slackness. Biodex dynamometer deformation has been estimated to account for up to 0.2 cm VL aponeurosis length change during maximal “isometric” torque production (Bojsen-Møller et al., 2003). Slack tendon refers to the idea that tendons become slack at short muscle-tendon complex length (Fukutani et al., 2017). During isometric torque production at 10% maximal force, VL fascicles shorten about six times more when the knee is extended than when flexed (about 3 cm versus 0.5 cm, Fukunaga et al., 1997). Since any non-zero muscle force may remove the slack in the tendon, considerable SEE length change may occur at small forces, requiring little work. In our experiment, a considerable portion of total fascicle shortening occurred at the smallest torque values (see Fig. 4). We found it challenging to estimate shortening work at very low torque levels, and dynamometer deformation and tendon slackness may have caused ours to be an overestimate of the work done on SEEs. However, such errors would tend toward a conservative, underestimate of the hypothesized force-rate cost.

Evidence for the proposed force-rate cost has previously been demonstrated in both muscle and whole-body experiments. For cyclic isometric contraction of mouse muscle *in situ* (1:3 on/off duty factor), rate of oxygen consumption is higher when cycle frequency is higher (Hogan et al., 1998). For prolonged isometric contraction of human VL muscle *in vivo*, rate of ATP utilization is higher when stimulation is cyclic (1:1 on/off duty factor) compared to constant (Chasiotis et al., 1987; Spriet et al., 1988). For cyclic isometric contraction of human calf muscle at constant frequency, metabolic cost decreases with increasing duty factor (Beck et al., 2020). While these studies have indicated a metabolic cost for cyclic muscle contraction, this cost was not parametrically related to muscle force-rate. We believe that force-rate, rather than work or force-time, could explain many such findings on metabolic cost of cyclic isometric contraction. In contrast to previous studies on isometric contraction varying on-off durations, we parametrically related such metabolic cost to muscle force-rate, which we find more suitable for the continuously varying forces of most daily movements. Furthermore, metabolic cost of cyclic movement has been observed to increase with movement frequency in leg swinging (Doke et al., 2005; Doke & Kuo, 2007), ankle bouncing (Dean & Kuo, 2011) and arm reaching (Wong et al., 2018). Altogether, the force-rate cost seems to be observable during a variety of movements and contraction conditions. We here provide evidence that the force-rate cost exists during voluntary isometric contraction, independently of work and force-time costs.

We have separated total metabolic cost into three terms: work-, force-time and force-rate. Such phenomenological division is potentially explained mechanistically as costs due to individual ATPases, particularly actomyosin (for work and force-time) and SERCA ATPases (for force-rate). Whether SERCA is primarily responsible for the force-rate cost could be tested in future experimental studies on isolated muscle, for example using cross-bridge (and therefore actomyosin ATPase) blocking agents (Barclay et al., 2007). We expect SERCA ATP consumption rate to increase with contraction frequency because of the increase in calcium release amplitudes, due to rate-limiting force production dynamics. Higher calcium amplitudes would require higher muscle activation levels, which agrees with the observed increase in quadriceps EMG amplitude with waveform frequency. Such properties are not included in Hill-type muscle models (Zajac, 1989), which do have activation dynamics attributed to calcium release (sometimes called ‘active state’) and muscle force-length and -velocity dependency, but not rate-limiting dynamics between calcium release and muscle force. As demonstration, we simulated the present experiment with a commonly used metabolic muscle model (OpenSim software, Uchida et al., 2016). It predicted a 2.4 W increase in metabolic rate with waveform frequency, only about 7% of the amount experimentally measured. Current models therefore appear to underestimate the energetic cost of cyclic force production. Improved estimates might be obtained by including rate-limiting dynamics, crudely modeled as a low-pass filter here, such as calcium release and binding (Baylor & Hollingworth, 2012) and Huxley-type crossbridge binding (van Soest et al., 2019). Future modelling studies could develop such mechanistic models that explain the force-rate cost.

Fast contraction of muscle may require high calcium release and pumping rates, at the cost of metabolic energy spend by SERCA. Such energetic penalty for rapid increase in muscle force may explain why (ground reaction) forces during human walking are smoother than expected based on work considerations (Rebula & Kuo, 2015). Nevertheless, there may be a considerable force-rate cost during human locomotion, as vastus lateralis fascicles produce relatively short bursts of high force (0 - 30% of stride; Bohm et al., 2018). The force-rate cost may also apply to a variety of other animals. In locust flight muscle, oxygen consumption during cyclic isometric contraction is 87% of the consumption during maximal power output, indicating high cost for cyclic muscle activation (Josephson & Stevenson, 1991). Locusts use synchronous muscle with one contraction per wing beat, whereas other insects such as beetles use asynchronous muscle with fewer contractions per beat (Josephson et al., 2001), which could reduce the activation cost as a potentially advantageous adaptation (Syme & Josephson, 2002). A force-rate cost could be an important factor in energy expenditure in a wide range of movements and species.

## Conclusion

We observed an increase in average net metabolic rate with frequency of cyclic torque production, which could not be explained by force-time nor by muscle fibre mechanical work. Average net metabolic rate was related to force-rate, suggesting that increasing muscle force abruptly requires more metabolic energy than when done slowly. We propose that this force-rate metabolic cost may be explained by an increase in the amount of required active calcium transport in the muscle fibre, and may be relevant for human and animal movement.

## Acknowledgements

The authors would like to thank Ashna Subramanium for help with data collection.

## Data availability

Ultrasound algorithm and typical example of ultrasound data is available on: https://github.com/timvanderzee/ultrasound-automated-algorithm

## References

Abbott, B. C., Bigland, B., & Ritchie, J. M. (1952). The physiological cost of negative work. The Journal of Physiology, 117(3), 380–390.

Barclay, C. J. (2015). Energetics of contraction. Comprehensive Physiology, 5(2), 961–995. https://doi.org/10.1002/cphy.c140038

Barclay, C. J., Constable, J. K., & Gibbs, C. L. (1993). Energetics of fast- and slow-twitch muscles of the mouse. The Journal of Physiology, 472(1), 61–80. https://doi.org/10.1113/jphysiol.1993.sp019937

Barclay, C. J., Woledge, R. C., & Curtin, N. A. (2007). Energy turnover for Ca2+ cycling in skeletal muscle. Journal of Muscle Research and Cell Motility, 28(4–5), 259–274. https://doi.org/10.1007/s10974-007-9116-7

Baylor, S. M., & Hollingworth, S. (2012). Intracellular calcium movements during excitation–contraction coupling in mammalian slow-twitch and fast-twitch muscle fibers. The Journal of General Physiology, 139(4), 261–272. https://doi.org/10.1085/jgp.201210773

Beck, O. N., Gosyne, J., Franz, J. R., & Sawicki, G. S. (2020). Cyclically producing the same average muscle-tendon force with a smaller duty increases metabolic rate. Proceedings of the Royal Society B: Biological Sciences, 287(1933), 20200431. https://doi.org/10.1098/rspb.2020.0431

Bergström, M., & Hultman, E. (1988). Energy cost and fatigue during intermittent electrical stimulation of human skeletal muscle. Journal of Applied Physiology (Bethesda, Md.: 1985), 65(4), 1500–1505. https://doi.org/10.1152/jappl.1988.65.4.1500

Bohm, S., Marzilger, R., Mersmann, F., Santuz, A., & Arampatzis, A. (2018). Operating length and velocity of human vastus lateralis muscle during walking and running. Scientific Reports, 8(1), 5066. https://doi.org/10.1038/s41598-018-23376-5

Bojsen-Møller, J., Hansen, P., Aagaard, P., Kjaer, M., & Magnusson, S. P. (2003). Measuring mechanical properties of the vastus lateralis tendon-aponeurosis complex in vivo by ultrasound imaging. Scandinavian Journal of Medicine & Science in Sports, 13(4), 259–265. https://doi.org/10.1034/j.1600-0838.2003.00301.x

Brenner, B., & Eisenberg, E. (1986). Rate of force generation in muscle: Correlation with actomyosin ATPase activity in solution. Proceedings of the National Academy of Sciences of the United States of America, 83(10), 3542–3546. https://doi.org/10.1073/pnas.83.10.3542

Chasiotis, D., Bergström, M., & Hultman, E. (1987). ATP utilization and force during intermittent and continuous muscle contractions. Journal of Applied Physiology (Bethesda, Md.: 1985), 63(1), 167–174. https://doi.org/10.1152/jappl.1987.63.1.167

Crow, M. T., & Kushmerick, M. J. (1982). Chemical energetics of slow- and fast-twitch muscles of the mouse. The Journal of General Physiology, 79(1), 147–166. https://doi.org/10.1085/jgp.79.1.147

Dean, J. C., & Kuo, A. D. (2011). Energetic costs of producing muscle work and force in a cyclical human bouncing task. Journal of Applied Physiology, 110(4), 873–880. https://doi.org/10.1152/japplphysiol.00505.2010

Doke, J., Donelan, J. M., & Kuo, A. D. (2005). Mechanics and energetics of swinging the human leg. The Journal of Experimental Biology, 208(Pt 3), 439–445. https://doi.org/10.1242/jeb.01408

Doke, J., & Kuo, A. D. (2007). Energetic cost of producing cyclic muscle force, rather than work, to swing the human leg. The Journal of Experimental Biology, 210(Pt 13), 2390–2398. https://doi.org/10.1242/jeb.02782

Ebashi, S., & Endo, M. (1968). Calcium and muscle contraction. Progress in Biophysics and Molecular Biology, 18, 123–183. https://doi.org/10.1016/0079-6107(68)90023-0

Fukunaga, T., Ichinose, Y., Ito, M., Kawakami, Y., & Fukashiro, S. (1997). Determination of fascicle length and pennation in a contracting human muscle in vivo. Journal of Applied Physiology (Bethesda, Md.: 1985), 82(1), 354–358. https://doi.org/10.1152/jappl.1997.82.1.354

Fukunaga, T., Kubo, K., Kawakami, Y., Fukashiro, S., Kanehisa, H., & Maganaris, C. N. (2001). In vivo behaviour of human muscle tendon during walking. Proceedings. Biological Sciences / The Royal Society, 268(1464), 229–233. https://doi.org/10.1098/rspb.2000.1361

Fukutani, A., Misaki, J., & Isaka, T. (2017). Relationship between joint torque and muscle fascicle shortening at various joint angles and intensities in the plantar flexors. Scientific Reports, 7(1), 290. https://doi.org/10.1038/s41598-017-00485-1

Hof, A. L. (1984). EMG and muscle force: An introduction. Human Movement Science, 3(1), 119–153. https://doi.org/10.1016/0167-9457(84)90008-3

Hogan, M. C., Ingham, E., & Kurdak, S. S. (1998). Contraction duration affects metabolic energy cost and fatigue in skeletal muscle. The American Journal of Physiology, 274(3), E397–402. https://doi.org/10.1152/ajpendo.1998.274.3.E397

Huang, H. J., Kram, R., & Ahmed, A. A. (2012). Reduction of metabolic cost during motor learning of arm reaching dynamics. Journal of Neuroscience, 32(6), 2182–2190.

Huang, N. E., Shen, Z., Long, S. R., Wu, M. C., Shih, H. H., Zheng, Q., Yen, N.-C., Tung, C. C., & Liu, H. H. (1998). The empirical mode decomposition and the Hilbert spectrum for nonlinear and non-stationary time series analysis. Proceedings of the Royal Society of London. Series A: Mathematical, Physical and Engineering Sciences, 454(1971), 903–995. https://doi.org/10.1098/rspa.1998.0193

Ichinose, Y., Kawakami, Y., Ito, M., & Fukunaga, T. (1997). Estimation of active force-length characteristics of human vastus lateralis muscle. Acta Anatomica, 159(2–3), 78–83. https://doi.org/10.1159/000147969

Inoue, M., Sakuta, N., Watanabe, S., Zhang, Y., Yoshikaie, K., Tanaka, Y., Ushioda, R., Kato, Y., Takagi, J., Tsukazaki, T., Nagata, K., & Inaba, K. (2019). Structural Basis of Sarco/Endoplasmic Reticulum Ca2+-ATPase 2b Regulation via Transmembrane Helix Interplay. Cell Reports, 27(4), 1221-1230.e3. https://doi.org/10.1016/j.celrep.2019.03.106

Josephson, R. K., Malamud, J. G., & Stokes, D. R. (2001). The efficiency of an asynchronous flight muscle from a beetle. Journal of Experimental Biology, 204(23), 4125–4139.

Josephson, R. K., & Stevenson, R. D. (1991). The efficiency of a flight muscle from the locust Schistocerca americana. The Journal of Physiology, 442, 413–429.

Kulig, K., Andrews, J. G., & Hay, J. G. (1984). Human Strength Curves. Exercise and Sport Sciences Reviews, 12(1), 417–466.

La Delfa, N. J., Sutherland, C. A., & Potvin, J. R. (2014). EMG processing to interpret a neural tension-limiting mechanism with fatigue. Muscle & Nerve, 50(3), 384–392. https://doi.org/10.1002/mus.24158

Lichtwark, G. A., & Wilson, A. M. (2008). Optimal muscle fascicle length and tendon stiffness for maximising gastrocnemius efficiency during human walking and running. Journal of Theoretical Biology, 252(4), 662–673. https://doi.org/10.1016/j.jtbi.2008.01.018

Lusk, G. (1909). The elements of the science of nutrition. Philadelphia, London, W. B. Saunders. http://archive.org/details/elementsscience03luskgoog

Margaria, R. (1968). Positive and negative work performances and their efficiencies in human locomotion. Internationale Zeitschrift Für Angewandte Physiologie Einschließlich Arbeitsphysiologie, 25(4), 339–351. https://doi.org/10.1007/BF00699624

Margaria, R. (1976). Biomechanics and energetics of muscular exercise. Oxford, London.

Rebula, J. R., & Kuo, A. D. (2015). The Cost of Leg Forces in Bipedal Locomotion: A Simple Optimization Study. PLOS ONE, 10(2), e0117384. https://doi.org/10.1371/journal.pone.0117384

Ruiter, C. J., Didden, W. J., Jones, D. A., & Haan, A. D. (2000). The force-velocity relationship of human adductor pollicis muscle during stretch and the effects of fatigue. The Journal of Physiology, 526 Pt 3, 671–681. https://doi.org/10.1111/j.1469-7793.2000.00671.x

Spriet, L. L., Soderlund, K., & Hultman, E. (1988). Energy cost and metabolic regulation during intermittent and continuous tetanic contractions in human skeletal muscle. Canadian Journal of Physiology and Pharmacology, 66(2), 134–139. https://doi.org/10.1139/y88-024

Syme, D. A., & Josephson, R. K. (2002). How to Build Fast Muscles: Synchronous and Asynchronous Designs. Integrative and Comparative Biology, 42(4), 762–770. https://doi.org/10.1093/icb/42.4.762

Uchida, T. K., Hicks, J. L., Dembia, C. L., & Delp, S. L. (2016). Stretching Your Energetic Budget: How Tendon Compliance Affects the Metabolic Cost of Running. PLOS ONE, 11(3), e0150378. https://doi.org/10.1371/journal.pone.0150378

van der Zee, T. J., & Kuo, A. D. (2020). Fully automated algorithm estimates muscle fascicle length from ultrasound image. BioRxiv, 2020.08.23.263574. https://doi.org/10.1101/2020.08.23.263574

van Ingen Schenau, G. J., Bobbert, M. F., & de Haan, A. (1997). Does Elastic Energy Enhance Work and Efficiency in the Stretch-Shortening Cycle? Journal of Applied Biomechanics, 13(4), 389–415. https://doi.org/10.1123/jab.13.4.389

van Soest, A. J., & Bobbert, M. F. (1993). The contribution of muscle properties in the control of explosive movements. Biological Cybernetics, 69(3), 195–204. https://doi.org/10.1007/BF00198959

van Soest, A. J., Casius, L. J. R., & Lemaire, K. K. (2019). Huxley-type cross-bridge models in largeish-scale musculoskeletal models; an evaluation of computational cost. Journal of Biomechanics, 83, 43–48. https://doi.org/10.1016/j.jbiomech.2018.11.021

Wong, J. D., Cluff, T., & Kuo, A. D. (2018). There is an energetic cost to movement jerk in human reaching. Society for Neuroscience, Washington, D.C.

Zajac, F. E. (1989). Muscle and tendon: Properties, models, scaling, and application to biomechanics and motor control. Critical Reviews in Biomedical Engineering, 17(4), 359–411.

